# Gas seepage pockmark microbiomes suggest the presence of sedimentary coal seams in the Öxarfjörður graben of NE-Iceland

**DOI:** 10.1101/348011

**Authors:** Guðný Vala Þorsteinsdóttir, Anett Blischke, M. Auður Sigurbjörnsdóttir, Finnbogi Óskarsson, Þórarinn Sveinn Arnarson, Kristinn P. Magnússon, Oddur Vilhelmsson

**Author notes:** (Passed away) (Þórarinn Sveinn Arnarson). Correspondence: Oddur Vilhelmsson.

## Abstract

Natural gas seepage pockmarks are found off and onshore in the Öxarfjörður graben, NE Iceland. The bacterial communities of two onshore seepage sites were analysed by amplicon sequencing of 16S rDNA, along with determining the geochemical characteristics, hydrocarbon content and the carbon isotope composition of the sites.

While one site was found to be characterised by biogenic origin of methane gas, with carbon isotope ratio δ^13^C [‰] = −63.2, high content of organic matter and complex hydrocarbons, the other site showed a mixed origin of the methane gas (δ^13^C [‰] = −26.6) with geothermal characteristics and lower organic matter content. While both sites harboured *Proteobacteria* as the most abundant bacterial phyla, the *Deltaproteobacteria* were more abundant at the geothermal site, and the *Alphaproteobacteria* at the biogenic site. The *Dehalococcoidia* class of the *Chloroflexi* phylum was abundant at the geothermal site while the *Anaerolineae* class was more abundant at the biogenic site. Bacterial strains from the seepage pockmarks were isolated on a variety of selective media targeting bacteria with bioremediation potential. A total of 106 strains were isolated and characterised, including representatives from the phyla *Proteobacteria, Bacterioidetes, Firmicutes*, and *Actinobacteria*. This article describes the first microbial study on gas seepage pockmarks in Iceland.

## Introduction

Natural gas seepage, the emission of gaseous hydrocarbons from the subsurface, has been studied extensively in the context of petroleum exploration because it can be used as an indicator of petroleum generation in subsurface sediments (1–3). Natural methane gas seepage is the result of subsurface generation or accumulation of methane and the methane concentration in the gas varies according to its source (4). At geothermal and hydrothermal sites, methane is generated by thermogenic processes and seeps up to the surface through cracks and pores, whereas, in deep sea sediments the accumulation of methane can result in cold seeps or methane hydrates where no direct input of heat is found. This is often linked to biogenic methane which is a product of microbial processes in various anaerobic environments, like bog lakes and sea sediments (5, 6). In many cases the methane generation is of mixed origin, that is both thermogenic and biogenic. For example, methane that is formed during early coalification processes (coal bed methane) is not only of thermogenic origin but also produced by microbes utilizing the lignite (7). In these environments one would expect to find microbes that participate in methanogenesis and are capable of methane oxidation, respectively.

Where natural methane gas seepage is active, pockmarks can develop that can be described as craters that are formed when gas or liquid comes seeping up from the subsurface (56). Gas seepage pockmarks can be regarded as hotspots for anaerobic oxidation of methane (AOM) that is often dependent on archaea and sulphate-reducing bacteria (47), but can in some cases be driven by bacteria through intra-aerobic-denitrification (9), or possibly reductive dehalogenation, as suggested in a recent study on an ice-covered Antarctic lake (10). Microbial communities of hydrocarbon gas seepage environments have been studied around the world, including the Gulf of Mexico (11), Pacific Ocean Margin (12), Cascadia Margin (13) and the Barents Sea (14), mainly because of their sulfate-reducing capabilities and AOM.

In Öxarfjörður bay, NE Iceland, natural gas seepage pockmarks are found both on the seafloor and on shore. Öxarfjörður is located along the lithospheric boundaries of the North-American and the Eurasian plates and forms a graben bounded by the Tjörnes Fracture Zone in the west and the eastern rim of the North Iceland Volcanic Zone in the east. Geothermal activity in Öxarfjörður bay is confined to three major fissure swarms, cross-sectioning the volcanic zone. The area is prevailed by the river delta of Jökulsá-á-Fjöllum, causing the Öxarfjörður bay to be even more dynamic in nature. Geological settings in the Öxarfjörður area were studied extensively in the 1990s (15–18), leading to the discovery that the methane-rich seepage gas likely originates from thermal alteration of lignite and coal seams from beneath the 1 km thick sediment (18). Taken together, these studies strongly suggest the presence of sedimentary lignite in the Öxarfjörður graben (19).

Very little geomicrobiological work has thus far been conducted in Iceland, with most environmental microbiology work being bioprospective in nature, often paying little attention to community structures or biogeochemical activity. Natural gas seeps such as those found in Öxarfjörður, have thus far not been investigated from a microbiological standpoint despite their unique character which makes them ideal for geomicrobiological studies as both sparsely vegetated geothermal gas seepage pockmarks and colder, more vegetated seeps are found in close proximity to one another. Each methane seep system is thought to be unique in terms of the composition of geological and biological features (8), so taking a snapshot of the microbial community at a methane gas seepage site can provide valuable insight into the dynamics of the system and initiate biological discoveries.

In this article, which has been available in preprint form (60), we report the first microbial analysis of the natural gas seepage pockmarks in Öxarfjörður. Hypothesizing that the microbial community in this environment ought to be dominated by methane oxidizing microbes, we performed microbial community analysis on 16S rRNA gene amplicon libraries from two sites differing in visible vegetation and complex hydrocarbon content. Further hypothesizing that these environments would be a source of hydrocarbon-degrading bacteria, we isolated a collection of microbes on various media and tested them for degradation of naphthalene.

## Materials and methods

### Sampling and in-field measurements

Samples were collected at Skógalón (site SX, 66°09’N, 16°37’W) on August 21st, 2014, and on September 11th, 2015, and at Skógakíll (site AEX, 66°10’N, 16°34’W) on August 13th, 2015 (Fig. 1). At site SX, where the natural gas seepage pockmarks are somewhat difficult to distinguish from ordinary marsh gas pockmarks, sites were selected where pockmarks were visibly active and appeared to form straight lines extending NW-SE. Temperature, pH and conductivity were measured *in-situ* during sampling with hand-held meters. Sediment samples were collected from shallow cores (50 cm) obtained using a corer constructed from a 3-cm diameter galvanized-iron pipe that was hammered into the ground using a sledgehammer, and transferred aseptically and mixed in sterile, airtight IsoJars (IsoTech laboratories, Champaign, Illinois), according to manufacturer’s protocol. Surface soil samples were collected aseptically directly into sterile IsoJars. Water samples were collected aseptically into sterile glass bottles. Gas samples were collected into evacuated double-port glass bottles by means of an inverted nylon funnel connected to silicone rubber tubing. All samples for microbial analysis were immediately put on dry ice where they were kept during transport to laboratory facilities at University of Akureyri where they were either processed immediately or stored in a freezer at −18°C until processing. Samples collected, along with *in-situ* measurements and types of sample are listed in Table 1.

**Fig. 1.**
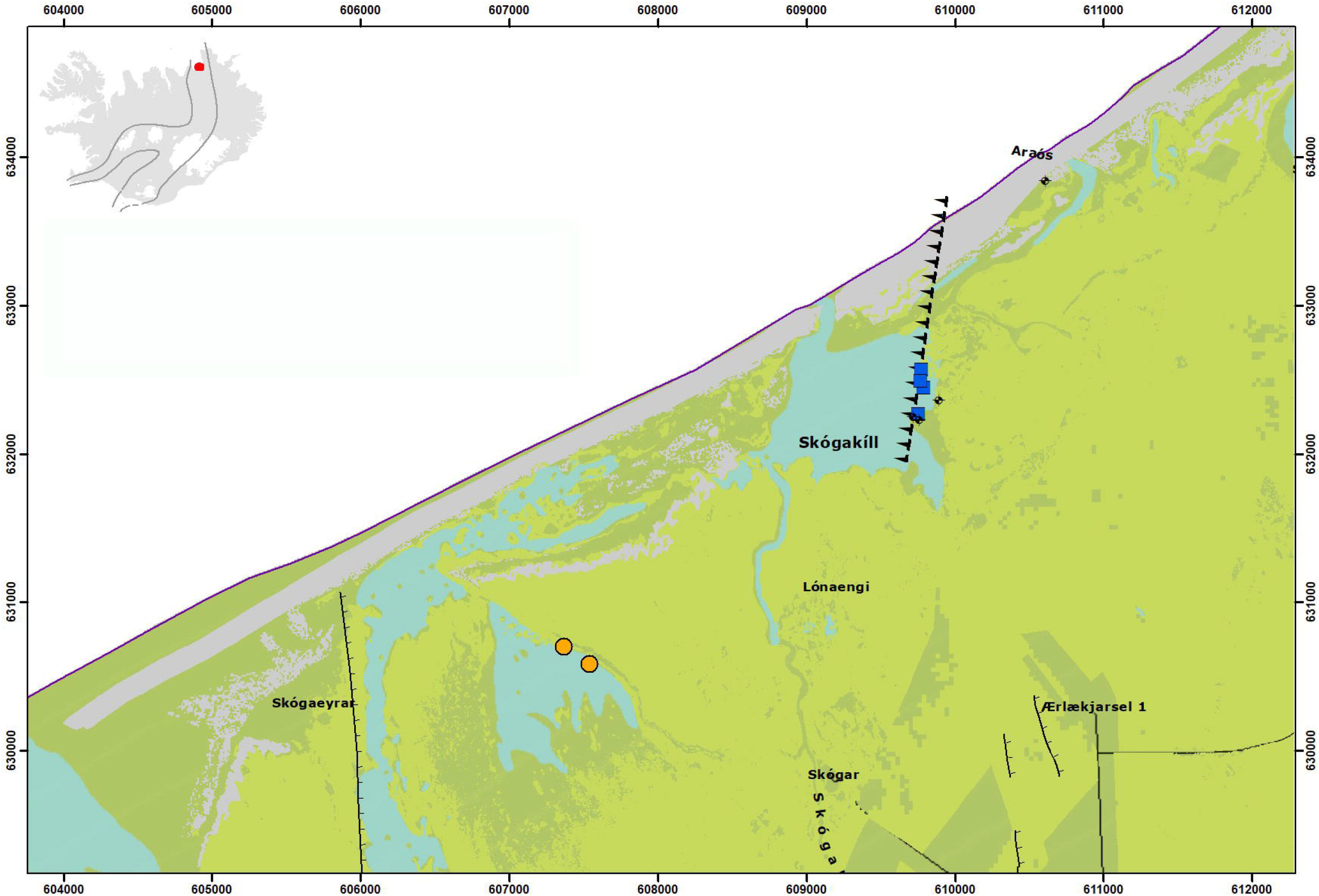
A map of the study area showing the AEX sampling sites (blue squares) and the SX sites (orange circles). Black diamonds indicate geothermal boreholes. Faults are inferred from the works of Sæmundsson et al. (31) and Ólafsson et al. (18). The insert shows the location of the study area in Iceland and the volcanic rift zone, bounded by the solid lines.

**Table 1.** *Location and description of sampling sites. Temperature and pH were measured* in situ *with a hand-held probe*.

| Site | Location | Coordinates | Description | pH | T (°C) |
| --- | --- | --- | --- | --- | --- |
| AEX-A | Skógakíll | 66°10'2.250 N<br>16°33'59.687 W | Gas seepage<br>sediment | 6,4 | 35,2 |
| AEX-B | Skógakíll | 66°10'8.014 N<br>16°33'56.422 W | Gas seepage<br>water | 6,4 | 53,3 |
| AEX-C | Skógakíll | 66°10'12.029 N<br>16°33'56.998 W | Mud-mire | ND | 30,4 |
| AEX-D | Skógakíll | 66°10'9.554 N<br>16°33'57.793 W | Sand-mud | 6,4 | 19,3 |
| SX-A | Skógalón | 66°09.222 N<br>016°37.089 W | Lagoon<br>sediment | 8,5 | 12,1 |
| SX-B | Skógalón | 66°09.244 N<br>016°37.240 W | Puddle water | 8,5 | 13,4 |

### Chemical analysis of geothermal fluids

Dissolved sulphide in the water samples was determined on-site by titration with mercuric acetate using dithizone in acetone as indicator (48). Major components in the water samples were determined at the laboratories of Iceland GeoSurvey (ÍSOR) in Reykjavík: Dissolved inorganic carbon was determined by alkalinity titration (pH 8.2 to 3.8), purging with nitrogen gas and back-titration (pH 3.8 to 8.2) as described previously (48). Silica was analysed by colorimetric determination of a silica-molybdate complex at 410 nm using a Jenway 6300 spectrophotometer. Total dissolved solids were determined by gravimetry. Anions were determined by suppressed ion chromatography on a ThermoScientific ICS-2100 with an AS-20 column. Major metals were analysed by atomic absorption spectrometry on a Perkin Elmer 1100B spectrometer. The composition of dry gas was also determined at the ÍSOR laboratories by gas chromatography on a Perkin Elmer Arnel 4019 light gas analyser equipped with HayeSep and MolSieve columns and three TCDs.

The concentration of trace elements in water samples were determined by ICP methods at the ALS Laboratories, Luleå, Sweden. Stable water isotopes (^2^H and ^18^O) were determined by mass spectrometry using a Delta V Advantage IRMS coupled with a Gasbench II at the Institute of Earth Sciences, University of Iceland.

**Headspace gas analysis from sediment samples** was performed at Applied Petroleum

Technologies, Kjeller, Norway, using standard techniques. Briefly as follows:

#### Sample preparation and extraction

Sediment samples were washed in water to remove mud before extraction using a Soxtec Tecator instrument. Thimbles were pre-extracted in dichloromethane with 7% (vol/vol) methanol, 10 min boiling and 20 min rinsing. The crushed sample was weighed accurately in the pre-extracted thimbles and boiled for 1 hour and rinsed for 2 hours in 80 cc of dichloromethane with 7% (vol/vol) methanol. Copper blades activated in concentrated hydrochloric acid were added to the extraction cups to cause free sulphur to react with the copper. An aliquot of 10% of the extract was transferred to a pre-weighed bottle and evaporated to dryness. The amount of extractable organic matter (EOM) was calculated from the weight of this 10% aliquot.

#### Deasphaltening

Extracts were evaporated almost to dryness before a small amount of dichloromethane (3 times the amount of EOM) was added. Pentane was added in excess (40 times the volume of EOM/oil and dichloromethane). The solution was stored for at least 12 hours in a dark place before the solution was filtered or centrifuged and the weight of the asphaltenes measured.

#### GC analysis of gas components

Aliquots of the samples were transferred to exetainers where 0.1-1ml were sampled using a Gerstel MPS2 autosampler and injected into a Agilent 7890 RGA GC equipped with Molsieve and Poraplot Q columns, a flame ionisation detector (FID) and 2 thermal conductivity detector (TCD). Hydrocarbons were measured by FID. H_2_, CO_2_, N_2_, and O_2_/Ar by TCD.

#### Carbon isotope analysis of hydrocarbon compounds and CO_2_

The carbon isotopic composition of the hydrocarbon gas components was determined by a GC-C-IRMS system. Aliquots were sampled with a syringe and analysed on a Trace GC2000, equipped with a Poraplot Q column, connected to a Delta plus XP IRMS. The components were burnt to CO_2_ and water in a 1000 °C furnace over Cu/Ni/Pt. The water was removed by Nafion membrane separation. Repeated analyses of standards indicate that the reproducibility of δ^13^C values is better than 1 ‰ PDB (2 sigma).

#### Carbon isotope analysis of low concentration methane using the Precon

The carbon isotopic composition of methane was determined by a Precon-IRMS system. Aliquots were sampled with a GCPal autosampler where CO_2_, CO and water were removed on chemical traps. Other hydrocarbons than CH_4_ and remaining traces of CO_2_ were removed by cryotrapping. The methane was burnt to CO_2_ and water in a 1000 °C furnace over Cu/Ni/Pt. The water was removed by Nafion membrane separation. The sample preparation system described (Precon) was connected to a Delta plus XP IRMS for δ ^13^C analysis. Repeated analyses of standards indicate that the reproducibility of δ^13^C values is better than 1 ‰ PDB (2 sigma).

#### GC of EOM fraction

A HP7890 A instrument was used. The column was a CP-Sil-5 CB-MS, length 30 m, i.d. 0.25 mm, film thickness 0.25 μm. C20D42 is used as an internal standard. Temperature programme: 50°C (1 min), −4 °C/min, −320 °C (25 min).

### Bacterial community analysis

Total DNA was extracted from 50 cm deep sediment samples in duplicates, using the PowerSoil kit (MoBio laboratories) following the manufacturer’s protocol. The DNA isolated was measured with Qubit fluorometer (Invitrogen, Carlsbad, CA) to confirm dsDNA in the samples. Four replicates from AEX1 and four from SX1 (Table 1.) were used to construct a paired-end library of the 16S rDNA hypervariable region V3/V4, using primers 337F and 805R, and sequenced on Illumina MiSeq platform by Macrogen, Netherlands. The data was processed and analysed using CLC Genomics Workbench 12.0 (https://www.qiagenbioinformatics.com/) and the CLC Microbial Genomics Module 4.0, with default parameters. Operational taxonomic units (OTU) were clustered by reference based OTU clustering using the SILVA 16S database v.132 for 97% similarity, allowing creation of new OTUs (*de novo*). For statistical analysis only alpha-diversity of samples was performed since the sequencing data only contained technical replicates, which does not allow analyses of beta-diversity. Differential abundance analysis (Likelihood Ratio test) was performed to see statistically significant differences in taxa between sampling sites.

### Initial culturing and isolation of bacteria

Samples were serially diluted to 10^-6^ in sterile Butterfield’s buffer and all dilutions plated in duplicate onto Reasoner’s agar 2A (Difco) and several selective and differential media including medium 9K for iron oxidizers (21), Mn medium for manganese oxidizers (990 mL basal agar B [0.42 g NaOAc, 0.1 g peptone, 0.1 g yeast extract, 15 agar, 990 mL sample water, autoclaved and cooled to 50°C], 10 mL pre-warmed filter-sterilized 1 M HEPES at pH 7.5, 100 μL filter-sterilized 100 mM Mn(II)SO_4_), Gui medium for laccase producers (990 mL basal agar B, 0,01% guiaicol), Hex medium for hexane degraders (990 mL basal agar B, 1.3 mL filter-sterilized 99 parts hexane/1 part dishwashing detergent), Naph medium for naphthalene degraders (basal agar B with several crystals of naphthalene added to the lid of inverted plates and then sprayed with fast blue for degradation indication), and 2,4-D medium for dichlorophenoxyacetate degraders (basal agar B supplemented with 2 mM 2,4-D). Three atmospheric incubation conditions were used: an unmodified atmosphere in sealed plate bags, a propane-enriched aerobic atmosphere in sealed plate bags flushed daily with propane, and an anaerobic, propane-supplemented atmosphere in anaerobic jars scrubbed of oxygen with a palladium catalyst (BBL GasPak) and monitored for anaerobicity with a resazurin strip. The jars were injected with 100 mL propane through a septum. Plates were incubated in the dark at 5, 15, or 22°C until no new colonies appeared (up to 4 weeks).

Colony morphotypes were examined by visual features, such as colour and form of elevation and margins. A representative of each morphotype was aseptically restreaked on fresh media and restreaked up to three times or until considered isolated strain. Stocks of isolates were prepared by suspending a loopful of growth in 1.0 mL 28% (v/v) glycerol and are stored at −70°C in the University of Akureyri culture collection.

### 16S rRNA gene-based identification of cultured strains

For each strain,1 μl of freezer stock was suspended in 25 μl of lysis buffer (1% Triton x-100, 20 mM Tris, 2 mM EDTA, pH 8,0) and incubated for 10 minutes at 95°C in the thermocyler (MJR PTC-200 thermocycler, MJ Research Inc. Massachusetts, USA). The lysis buffer solution (1 μl), or 1 μl of extracted DNA (using UltraClean® Microbial DNA Isolation Kit (MoBio Laboratories, Carlsbad, California, USA)), was used as a DNA template for Polymerase chain reaction (PCR) using Taq DNA-polymerase to amplify the DNA using the ‘universal’ bacterial primers 8F (5’-AGTTTGATCCTGGCTCAG’3) and 1522R: (5’-AAGGAGGTGATCCAGC CGCA-’3) (57).

The PCR reaction was conducted as follows: 95°C for 3 min, followed by 35 cycles of 95°C for 30 sec, 50°C for 30 sec and 68°C for 90 sec, then final extension at 68°C for 7 min. The PCR products were loaded on 0,8% agarose gel and run at 100 V for approximately 30 minutes to verify the presence of approximately 1500-bp amplicons.

The PCR products were purified for sequencing using 22 μl of sample in 10 μl ExoSap Mix (mix for 50 reactions; 1.25 μl Exonuclease I [20 U/μl], 2.5 μl Antarctic phosphatase [5 U/μl] (New England BioLabs Inc.), 496.5 μl distilled H_2_O). The products were incubated at 37°C for 30 minutes and heated to 95°C for 5 minutes.

The purified PCR products were sequenced with BigDye terminator kit on Applied Biosystem 3130XL DNA analyser (Applied Biosystems, Foster City, California, USA) at Macrogen Europe, Amsterdam, the Netherlands. Two sequencing reactions were run for each strain, using the primers 519F (5’-CAGCAGCCGCGGTAATAC-’3) (58) and 926R (5’-CCGTCAATTCCTTTGAG TTT-’3) (59). The resulting sequences were trimmed using ABI Sequence Scanner (Applied Biosystems), the forward sequence and the reverse complement of the reverse sequence aligned and combined, and taxonomic identities obtained using the EzTaxon server (22).

## Results

**Water chemical analysis** revealed several differences in major components at the two sites (Table 2). Although the pH of the water did not differ significantly as judged by a two-tailed Student’s t-test, electrical conductivity was nearly 12-fold higher at the AEX site, significant at the 99.5% confidence level. The AEX site water contained more than 420-times as much silica as did the SX site water, and several other ions, such as sodium, potassium chloride and bromide were also found to be present at significantly higher levels at the AEX site, underscoring the more geothermal character of the environment (Table 2).

**Table 2.** Major physicochemical characteristics^1^ of water from the two study sites.

|  | AEX | SX |  |
| --- | --- | --- | --- |
| pH at 21°C | 7.57 ± 0.31 | 8.64 ± 0.73 | (p = 0.19) |
| Conductivity at 25°C<br>(µScm<sup>-1</sup>) | 8945 ± 728 | 773 ± 392 | (p = 0.005) |
| DIC (as ppm CO<sub>2</sub>) | 44.8 ± 31.0 | 94.5 ± 43.1 | (p = 0.32) |
| SiO<sub>2</sub> (ppm) | 147.5 ± 2.1 | 0.35 ± 0.07 | (p = 0.0001) |
| Na (ppm) | 1900 ± 56.6 | 107.4 ± 57.5 | (p = 0.001) |
| K (ppm) | 104.9 ± 18.6 | 4.01 ± 1.96 | (p = 0.02) |
| Mg (ppm) | 24.0 ± 15.9 | 17.8 ± 10.5 | (p = 0.69) |
| Ca (ppm) | 311.5 ± 111.0 | 17.1 ± 9.0 | (p = 0.06) |
| F (ppm) | 0.45 ± 0.13 | 0.21 ± 0.01 | (p = 0.11) |
| Cl (ppm) | 3355 ± 276 | 169.4 ± 105.5 | (p = 0.004) |
| Br (ppm) | 12.2 ± 0.8 | 0.34 ± 0.31 | (p = 0.002) |
| SO<sub>4</sub> (ppm) | 215 ± 76.4 | 5.13 ± 0.60 | (p = 0.06) |

<sup>1</sup>Dissolved sulphide was determined on-site by titration with mercuric acetate using dithizone as indicator (48). Dissolved inorganic carbon was determined by alkalinity titration and back-titration (48). Silica was analysed by colorimetric determination of a silica-molybdate complex at 410 nm. Total dissolved solids were determined by gravimetry. Anions were determined by suppressed ion chromatography. Major metals were analysed by atomic absorption spectrometry.

**Headspace gas analysis** revealed similar amounts of hydrocarbon gas at the two sites (Table 3). Although methane content was found (n=3) to be lower at the AEX site than at the SX site, the difference was not deemed significant by a two-tailed Student’s t-test (not shown). Isotope composition suggests a thermogenic origin of the AEX-site headspace gas, whereas a biogenic origin is suggested for the SX-site headspace gas (Table 3). Thermogenic origin of the AEX gas was further supported by a high methane/ethane ratio (17.8). Both the composition of the headspace gas and the methane isotope composition were similar to those reported by Ólafsson *et al*. for borehole gasses in the Skógalón area (18). EOM fractions showed difference in hydrocarbon content in terms of lower-chain hydrocarbons in AEX and higher amounts of longer-chain hydrocarbons at the SX site (Fig. 2).

**Table 3.** Headspace gas analysis^1^ on sediment samples from seepage pockmarks at the two study sites. The concentrations are shown as parts of Total Hydrocarbon Gas (THCG).

|  | AEX | SX |
| --- | --- | --- |
| N<sub>2</sub> [%total] | 39.2 (±31.5) | 81.2 (±2.62) |
| O<sub>2</sub> + Ar [%total] | 8.46 (±8.53) | 14.10 (±5.66) |
| ppm THCG | 5.09×10<sup>5</sup> (±3.88×10<sup>5</sup>) | 0.389×10<sup>5</sup> (±0.296×10<sup>5</sup>) |
| CO<sub>2</sub> [% THCG] | 99.9 (±0.23) | 93.2 (±1.77) |
| Methane (CH<sub>4</sub>)<br>[% THCG] | 46.1×10<sup>-3</sup> (±48.3×10<sup>-3</sup>) | 6.78 (±1.66) |
| Ethane (C<sub>2</sub>H<sub>6</sub>)<br>[% THCG] | 2.60×10<sup>-3</sup> (±4.50×10<sup>-3</sup>) | 0 (±0) |
| Propane (C<sub>3</sub>H<sub>8</sub>)<br>[% THCG] | 1.03×10<sup>-3</sup> (±1.79×10<sup>-3</sup>) | 0 (±0) |
| iso-Pentane (*i*-C<sub>5</sub>H<sub>12</sub>)<br>[% THCG] | 18.7×10<sup>-3</sup> (±32.3×10<sup>-3</sup>) | 8.00×10<sup>-3</sup> (±11.3×10<sup>-3</sup>) |
| Pentane (*n*-C<sub>5</sub>H<sub>12</sub>)<br>[% THCG] | 73.7×10<sup>-3</sup> (±127×10<sup>-3</sup>) | 49.5×10<sup>-3</sup> (±70.0×10<sup>-3</sup>) |
| Benzene (C<sub>6</sub>H<sub>6</sub>)<br>[% THCG] | 3.80×10<sup>-3</sup> (±6.24×10<sup>-3</sup>) | 7.15×10<sup>-3</sup> (±5.45×10<sup>-3</sup>) |
| Methane δ<sup>13</sup>C [‰] | -26.6 | -63.2 |
| CO<sub>2</sub> δ<sup>13</sup>C [‰] | -3.3 | nd |

<sup>1</sup>Samples were extracted in dichloromethane/methanol, deasphalted in pentane, and analysed by gas chromatography.

**Fig. 2.**
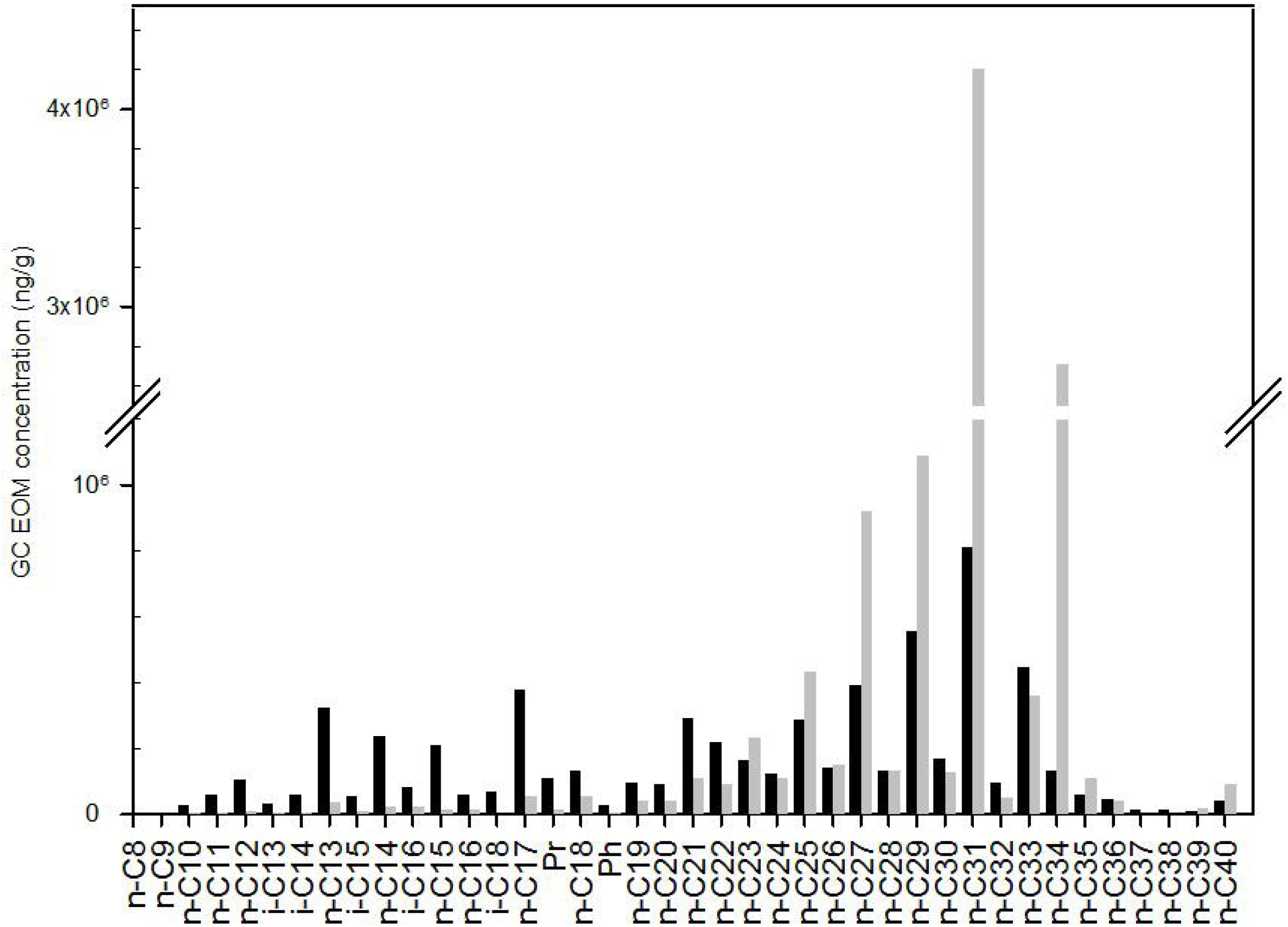
Hydrocarbon content in sediment samples from the natural gas seepage pockmarks. Extractable organic matter (EOM) concentration as determined by gas chromatography is compared between the AEX (dark columns) and SX (light columns) study sites (two samples per site). Error bars are omitted for clarity.

### Bacterial community results

Four technical replicates from each study site were used to compare the microbial communities in the gas seepage pockmarks of Skógalón (SX) and Skógakíll (AEX). The DNA extraction yielded on average 2.8 μg/ml (± 0,3 μg/ml) and the amplicon library generated on average 42.8 ng/μL (± 1,3 ng/μL) of amplicons with the length of 587±5 bp. Over 4 million paired sequences, were analysed and trimmed to the average of 523,241 reads per sample with the length of 301 bp. A total of 595,137 reads generated the OTU table after filtering out chimeric sequences, where predicted OTUs were in total 26,786 OTUs (Table 4).

**Table 4.**
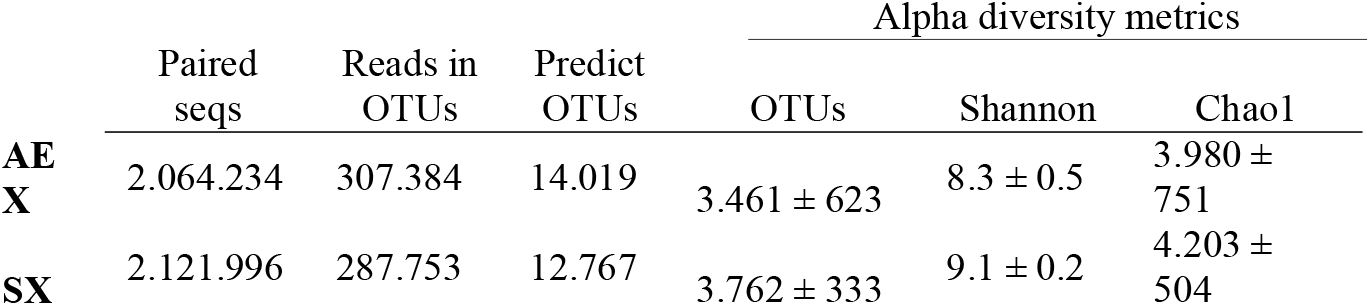
Number of predicted operational taxanomic units (OTUs) and alpha diversity metrics as calculated at 25,000 sequences from the two study sites.

Alpha diversity metrics were measured at the depth of 60,000 sequences per sample. Based on rarefaction analysis, the number of OTUs had reached plateau at 25,000 sequences, meaning the dataset was sufficient to estimate the diversity of the bacterial communities in the natural seepage pockmarks. The species richness as estimated with Chao1 index and Shannon’s diversity index are shown in Table 4.

### Taxonomic composition

The focus was set on analysing the most relative abundant taxa, so after filtering out chloroplast OTUs, the OTUs with the lowest combined abundance (<= 1% of total reads) were omitted. A total of 14 bacterial phyla was observed as the most abundant at AEX and SX sites, divided up to 23 classes and 45 observed genera (Fig. 3). The *Proteobacteria* phylum had the most abundant OTUs at both AEX and SX sites, 28% and 30%, respectively. At the AEX site, *Proteobacteria* was followed by *Chloroflexi* (22%) and *Aminicenantes* (10%) at phylum level. At the SX site, the *Bacteroidetes* had significantly higher abundance than in the AEX site, with relative abundance of 24%, followed by *Chloroflexi* (13%).

The bacterial community structure differed between sites, but only one class, within the phylum of *Aminicenantes* was found by likelihood ratio analysis to be significantly more abundant at the AEX site than at the SX site. An unnamed order within the *Bacteriodetes* was found to be significantly more abundant at the SX site compared to the AEX. The family of *Syntrophaceae* within the class of *Deltaproteobacteria* was more abundant at the SX site with 2.6-fold higher relative abundance. OTUs of three genera within the *Clostridia* class had over 3.0-fold higher relative abundance at AEX than SX site.

**Fig. 3.**
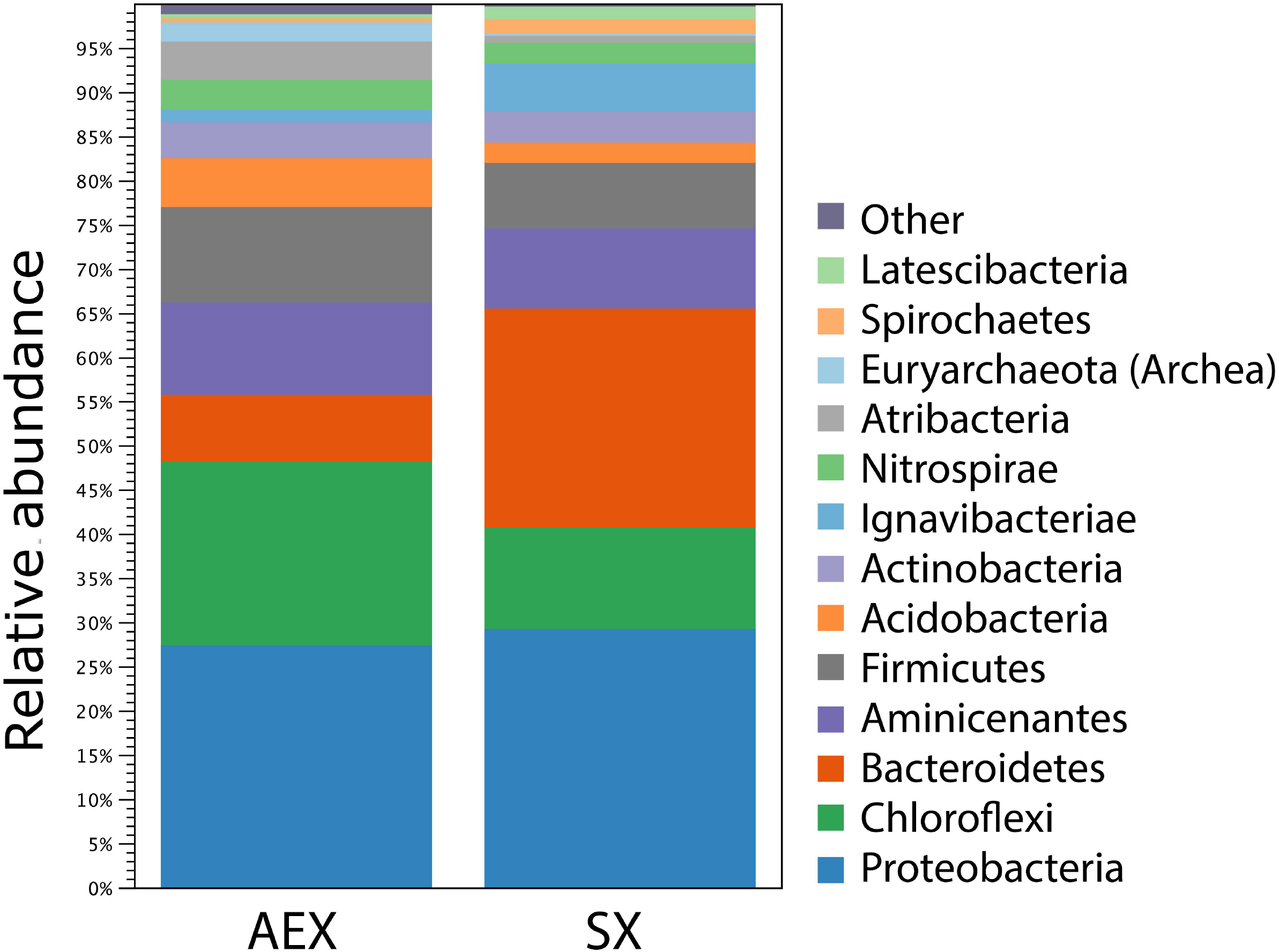
Bacterial community structure in natural gas seepage pockmarks at Skógakíll (AEX) and Skógalón (SX) sites, presented as the relative abundance of bacterial phyla from amplicon sequencing of V3-V4 in 16S rDNA. Operational taxanomic units (OTUs) with relative abundance lower than 0.1 were omitted.

### Cultured microbiota and isolates

Plate counts after 7 days at 22°C (Table 5) of samples from site SX indicated the presence of substantial communities of naphthalene and hexane degraders, particularly under aerobic conditions. One hundred and eighty-six colonies were restreaked for isolation in pure culture (Table 9 in supplements).

**Table 5.**
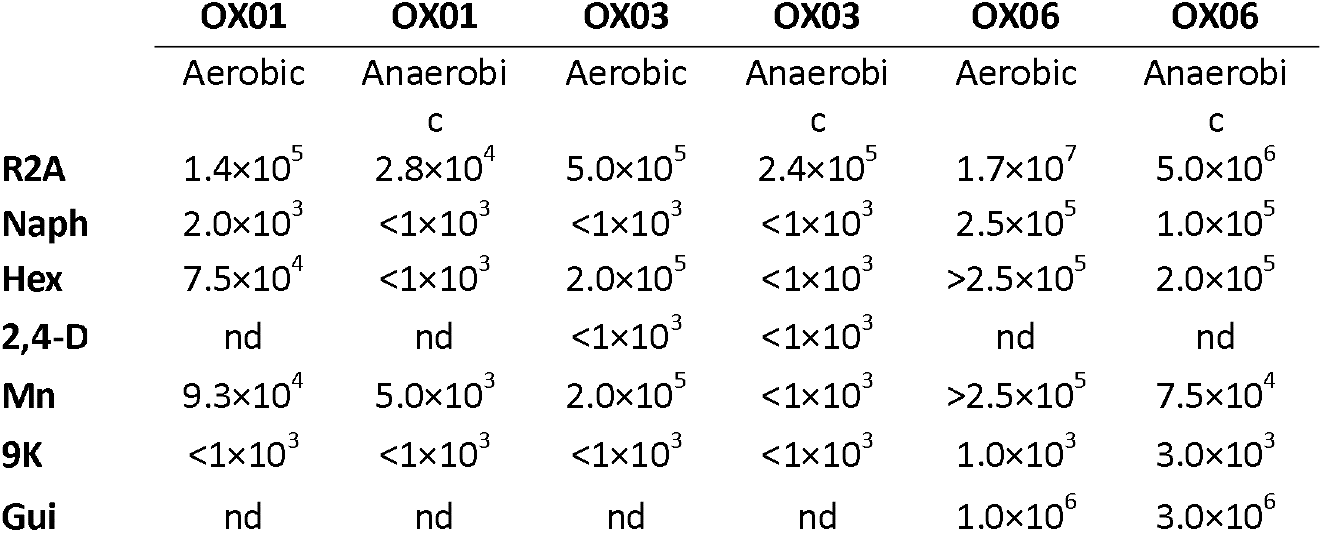
Colony-forming units per gram sediment sample after 7 days at 22 °C on selective media.

Putatively facultative chemoautotrophs were surprisingly numerous judging by growth on Mn media, but the extremely restrictive medium 9K only yielded a few colonies, all from sample OX06. Spraying colonies with fast blue confirmed the presence of alpha-naphthol in some of the colonies on Naph-agar, but not all. Strains OX0102 and OX0103 tested positive for naphthalene degradation by fast blue; strains OX0304 and OX0306 tested positive for 2,4-dichlorophenoxy acetic acid degradation by fast blue. One hundred and six strains have been identified by partial 16S rDNA sequencing using the Sanger method and found to comprise 38 genera in 8 classes (Table 6, Table 10 in supplements).

**Table 6.**
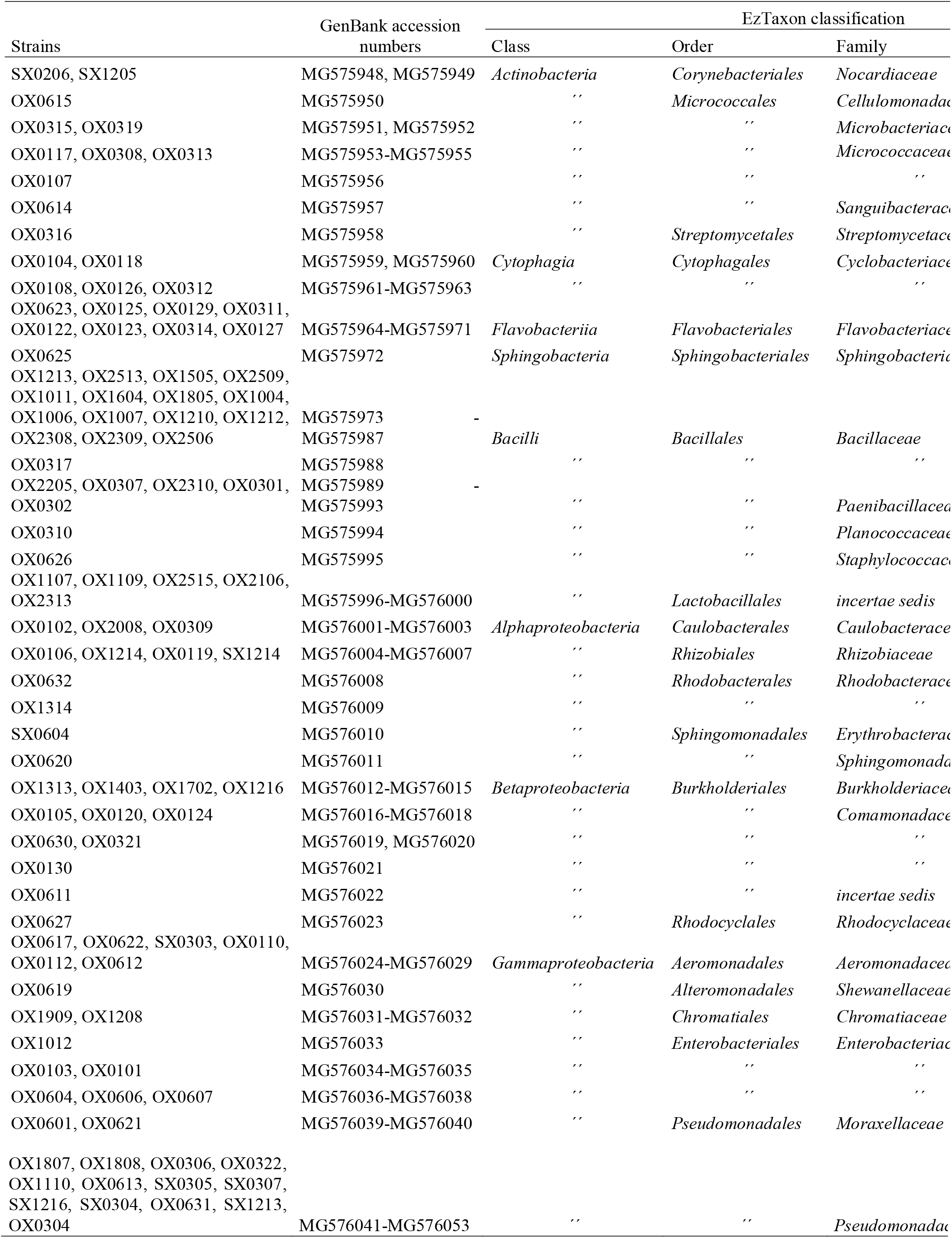
Bacterial isolates and their taxonomic classification by partial 16S rRNA gene sequencing.

## Discussion

The study sites, AEX and SX, were found to be distinct in terms of geochemistry. The AEX site contained higher concentrations of silica, very similar to the concentrations of previous studies on geothermal activity in Öxarfjörður (18) indicating geothermal water coming from the pockmarks. The sodium chloride concentration was higher than in previous studies which implies a mixture of seawater with the geothermal water. However, the pockmarks at AEX are located in a river delta and water samples were taken at the pockmark surface so the intermixture of seawater is not surprising. The water chemistry at the SX site shows low concentration of silica indicating little or no geothermal activity and less intermixture of seawater than at AEX. The stable isotope ratio δ^13^C of methane also indicates a biogenic origin of the methane at SX, while at AEX the δ^13^C suggests a mixture of thermogenic and biogenic origin of methane, which can most likely be linked to microbial lignite utilization at the site as well as the geothermal activity previously described (18). The hydrocarbon content also shows more complex and longer-chain hydrocarbons at SX. This can be related to more vegetation and organic matter accumulation, in contrast with lower-chain hydrocarbons at the AEX site with less vegetation (Fig.2). These geochemical factors underline how disparate the two sites are: the AEX pockmarks containing geothermal groundwater with thermogenic methane generation and the SX pockmarks the result of biogenic natural gas accumulation. The location of the SX site and the lining up of the pockmarks can easily suggest thermogenic methane seepage at the site, but our analysis of the SX-site pockmarks explored in this study is more suggestive of marsh gas seepage, whereas at the AEX site, a thermogenic origin is more strongly supported. The difference in hydrocarbon content of the two sites, is most likely to explain the difference in biodiversity presented with biodiversity indices in Table 4. Howewer, further studies are needed to demonstrate the correlation between hydrocarbon content and biodiversity.

The seepage pockmarks were found to harbour diverse microbiotas consisting largely of anaerobic heterotrophs. Given the lack of visible vegetation at the AEX site, available organic matter seems likely to be restricted to the gas seep itself, to a large extent. This kind of environment thus contains a microbial community composed largely of facultative chemolithotrophs and oxidizers of methane, lighter alkanes, and aromatics. The inter-site diversity of both sampling sites was notable. However, several groups of bacteria were shown to vary in relative abundance between the two sampling sites, as discussed below.

### Hydrocarbon and methyl halide metabolism

The high relative abundance of *Dehalococcoidia* in the microbial consortia at the study sites, particularly site AEX, is noteworthy and underscores the profound effect that petrochemical seepage has on the composition of the local microbiota. The class *Dehalococcoidia* contains at the present time only one validly described order (*Dehalococcoidales*), one family (*Dehalococcoidaceae*), and three genera (*Dehalococcoides, Dehalobium* and *Dehalogenimonas*), comprising a total of four species, all of which are capable of anaerobic reductive dehalogenation (23–27).

A large fraction of the dehalococcoidal OTUs in this study were found to be assignable to the order *Dehalococcoidales* (Table 7), where the majority (9120 of 9579) being further assignable to the family *Dehalococcoidaceae*, containing the described, dehalorespiring members of this class. In further support of dehalorespiration being an important process in the seepage pockmark microbiotas, genera known to contain facultative dehalorespirers, like the betaproteobacterial genus *Dechloromonas* (28) and the deltaproteobacterial genus *Anaeromyxobacter* (29), were significantly more abundant at the AEX site. Furthermore, cultured bacteria from the Öxfjörður seeps, while not including *Dehalococcoidia*, do include isolates assigned to genera known to include aerobic facultative dechlorinators, such as *Dechloromonas* and *Shewanella* (Table 7).

**Table 7.** *Fractional abundance of* Chloroflexi *classes and orders in the seepage pockmarks microbiomes as determined with amplicon sequencing*.

| Class | AEX<sub>1</sub> | SX<sup>1</sup> | Order | AEX<sub>2</sub> | SX<sup>2</sup> |
| --- | --- | --- | --- | --- | --- |
| *Anaerolineae* | **0.65** | **0.87** |  |  |  |
|  |  |  | *Anaerolineales* | 0.62 | 0.94 |
|  |  |  | *Caldilineales* | 0.02 | 0.00 |
|  |  |  | ‘MSB-5B2’ | **0.07** | **0.00** |
|  |  |  | ‘SBR1031’ | **0.08** | **0.01** |
|  |  |  | ‘RBG-13-54-9’ | 0.02 | 0.01 |
|  |  |  | Others | **0.19** | **0.04** |
| *Dehalococcoidia* | **0.30** | **0.08** |  |  |  |
|  |  |  | *Dehalococcoidales* | 0.02 | 0.22 |
|  |  |  | ‘GIF9’ | **0.70** | **0.26** |
|  |  |  | ‘FS117-23B-02’ | **0.06** | **0.02** |
|  |  |  | Others | 0.22 | 0.50 |
| ‘KD4-96’ | 0.02 | 0.04 |  |  |  |
| Others | 0.03 | 0.01 |  |  |  |

<sup>1</sup>Fraction of *Chloroflexi* paired-end sequences (total of 86,729 paired-end reads). Figures in **bold** indicate a significant difference between the AEX and SX microbiota by a two-tailed two-sample Student’s *t*-test ( $p < 0.05$ ).
<sup>2</sup>Fraction of class-level paired-end sequences (total of 44,384 paired-end *Anaerolineae* reads and 36,549 paired-end *Dehalococcoidia* reads). Figures in **bold** indicate a significant difference between the AEX and SX microbiota by a two-tailed two-sample Student’s *t*-test ( $p < 0.05$ ).

The as-yet unnamed and uncharacterized order GIF9 was highly abundant in the AEX and SX microbiomes (Table 7) and may consist of bacteria that possess other metabolic pathways than just organohalide respiration. The order is suggested to be an important bacterial group for the degradation of organic matter in sediments (49). Thus, a recent metagenomic study indicated that some members of this group may be homoacetogenic fermenters that possess a complete Wood-Ljungdahl CO_2_ reduction pathway (31). It should thus be stressed that the presence of a large contingent of *Dehalococcoidia*, as was found to be the case in the present study, need not necessarily be indicative of dehlaorespiration constituting a major metabolic activity in the environment under study. Indeed, considerable variation in metabolic characteristics occurs in most well-characterized bacterial classes and hence it must be considered likely that other, perhaps non-dehalorespiring taxa remain to be characterized within this class.

Recently, it was suggested, in part because of the notable abundance of *Dehalococcoides*, that in certain Antarctic lakebeds, anaerobic methane oxidation may be fuelled by reductive dehalogenation (10). The results of the present study are suggestive of the presence of such an ecosystem in the methane seeps in the Öxarfjörður graben. Methyl halides are often associated with coal combustion (32), further suggesting subsurface interaction of geothermal matter with lignite as a source of chloromethane.

Also among abundant groups observed in the present study, the *Atribacteria* (group ‘OP9’) are often found to be predominant in methane-rich anaerobic environments such as marine sediments and subseafloor “mud volcanoes” (43, 44). Although they have not been directly linked to AOM in these environments, they have been suggested to mediate AOM in some cold seep environments (10). In general, the *Atribacteria* are thought to play heterotrophic roles, likely fermentative (43, 45), but a single-cell genomics study on representative *Atribacteria* suggests that these organisms may be indirectly responsible for methane production through the production of acetate or CO_2_ (43, 46).

Another highly abundant class within the *Chloroflexi* was the *Anaerolineae* (Table 7), a class originally described as consisting of strictly anaerobic chemo-organotrophs (33), and frequently detected in subsurface environments (34–37). However, due to the scarcity of cultured representatives, the metabolic capabilities of this class have remained elusive. A recent study of seven single-cell genomes from deep submarine hydrothermal vent sediments indicated the presence of a Wood-Ljungdahl CO_2_ reduction pathway, as well as a number of ABC transporters, and in one case a putative reductive dehalogenase (38). In the present study, the *Anaerolineae* appear fairly diverse, with 81% of *Chloroflexi* paired-end reads being assigned to four orders: the *Anaerolineales* and three putative orders without cultured representation, envOPS12, GCA004, and SHA-20 (Table 7).

### *Proteobacteria* and the sulfur cycle

Members of the *Proteobacteria* phylum observed in the seepage pockmarks mainly consisted of *Deltaproteobacteria* and *Alphaproteobacteria* (Table 8). The alphaproteobacterial fraction was fairly homogeneous, consisting mostly of reads assigned to the order *Rhizobacteriales*, of which 77% could be assigned to a single genus, *Bosea*, a genus of chemolithoheterotrophs noted for their ability to oxidize inorganic sulfur compounds (39). The deltaproteobacterial fraction was found to be more diverse although most of the OTUs could be assigned to either of two orders, the *Syntrophobacterales* and the *Desulfobacterales* (Table 8), both of which contain mostly, albeit not exclusively, sulfatereducing organisms.

**Table 8.** *Fractional abundance of* Proteobacteria *classes and orders in the seepage pockmarks microbiomes as determined with amplicon sequencing*.

| Class | AEX<sup>1</sup> | SX<sup>1</sup> | Order | AEX<sup>2</sup> | SX<sup>2</sup> |
| --- | --- | --- | --- | --- | --- |
| *Delta-proteobacteria* | **0.51** | **0.23** |  |  |  |
|  |  |  | *Syntrophobacterales* | **0.53** | **0.70** |
|  |  |  | *Desulfobacterales* | 0.08 | 0.00 |
|  |  |  | *Myxococcales* | 0.02 | 0.02 |
|  |  |  | *Desulfarculales* | **0.07** | **0.00** |
|  |  |  | *Desulfuromonadales* | 0.01 | 0.01 |
|  |  |  | 'Sva0485' | 0.18 | 0.16 |
|  |  |  | Others | 0.11 | 0.11 |
| *Alpha-proteobacteria* | **0.22** | **0.10** |  |  |  |
|  |  |  | *Rhizobiales* | **0.76** | **0.96** |
|  |  |  | *Rhodobacterales* | 0.09 | 0.01 |
|  |  |  | Others | 0.15 | 0.03 |
| *Gamma-proteobacteria* | **0.27** | **0.08** |  |  |  |
|  |  |  | *Betaproteobacteriales* | 0.35 | 0.49 |
|  |  |  | Other | 0.65 | 0.51 |

<sup>1</sup>Fraction of *Proteobacteria* paired-end sequences (total of 152,153 paired-end reads). Figures in **bold** indicate a significant difference between the AEX and SX microbiota by a two-tailed two-sample Student's *t*-test ( $p < 0.05$ ).
<sup>2</sup>Fraction of class-level paired-end sequences (total of 83,764 paired-end *Deltaproteobacteria* reads and 43,823 paired-end *Alphaproteobacteria* reads). Figures in **bold** indicate a significant difference between the AEX and SX microbiota by a two-tailed two-sample Student's *t*-test ( $p < 0.05$ ).

The *Syntrophobacterales*, known to be frequently associated with anoxic aquatic environments (40), are significantly enriched in the SX marshland site as compared to the AEX site, perhaps reflecting an influx of marshland-associated bacteria into the seepage pockmark environment. Most of the *Syntrophomonadales* reads (77%) can be assigned to the family *Syntrophaceae*, which contains both sulfate-reducing and non-sulfate-reducing bacteria (41). Many of the *Syntrophaceae* reads (59%) could not be confidently assigned to genera, rendering the question of the importance of sulfate reduction of this taxon in the seepage pockmarks unresolved. However, taken together with the high abundance of the sulfate-reducing order *Desulfobacterales*, sulfate reduction is likely to be a major process in the seepage pockmarks, likely supporting AOM consortia. Families within *Desulfobacterales* have been reported to actively oxidize short and long chain alkanes and are suggested to be the key alkane degraders in marine seeps (42). Furthermore, considering the high abundance of the sulfur-oxidizing *Bosea*, we can surmise that *Proteobacteria* consitute an important driver of sulfur cycling within the seepage pockmark microbiota.

### Bioremediative potential of isolated bacterial strains

Microbial communities containing facultative chemolithotrophs and hydrocarbon oxidizers, could be valuable for bioremediation of petroleum contamination in basaltic oligotrophic environments (54, 55) such as Icelandic beach environments. Previously it has been demonstrated that the microbial activity in Icelandic soils is more affected by substrate availability than temperature (50, 51), wich means the possibility of bioremediative strategies like biostimulation is quite possible. Screening environmental isolates for possible bioremediative capabilities is therefore meaningful for further research into potential bioremediation in contaminated seashore environments in Iceland. Two of the isolated strains showed degradation potential of naphthalene. Strain OX0102 was assigned to the genus *Brevundimonas* (Table 10 in supplements) and Strain OX0103 was most closely related to bacteria in the genus *Rahnella* (Table 10 in supplements), but strains from both genera have been isolated with high hydrocarbonoclastic activity in relation to the degradation of naphthalene (52, 53). Strains OX0304 and OX0306, both assigned by EzTaxon as *Pseudomonas* species (Table 10 in supplements), showed putative degradation of 2,4-dichlorophenoxy acetic acid. Furthermore, strain OX0627 was assigned as a member of *Dechloromonas*, but isolates from that genus are capable of anaerobic oxidation of benzene (30) and could possibly be used for bioremediation, as well as the isolated strains mentioned above. This substantiates the significance of isolated strains presented in the study for future research in Iceland, related to hydrocarbon degradation and bioremediation.

## Concluding remarks

This study presents the first microbial community analysis of a gas seepage site in Iceland and compares the microbiomes of neighboring biogenic and thermogenic gas seepage pockmarks in the Jökulsá-á-Fjöllum river delta. It thus provides insights into microbial communities in these unusual environments and raises compelling questions on the connection between the gas origin and the pockmark microbiota, establishing the need for further geomicrobiological research in Icelandic natural gas seeps. The microbial communities associated with the pockmarks show higher biodiversity in the biogenic gas seep than in the thermogenic one, presenting a diverse microbiota that consists largely of anaerobic heterotrophs. The abundant taxa in the pockmarks indicate that the microbial community is most likely involved in hydrocarbon degradation linked to sulfur cycling and AOM, and the abundance of *Dehalococcoidia* suggests the presence of anaerobic reductive dehalogenation in natural gas seepage pockmarks of thermogenic origin. Several strains isolated in this study have demonstrable hydrocarbon-degrading activity, and thus comprise a pivotal resource for future studies in bioremediation and enviromental biotechnology in Iceland.

## Acknowledgments

This work was funded by Orkustofnun and Orkurannsoknarsjodur Landsvirkjunar. We thank Geir Hansen & Co. at Applied Petroleum Technology for their contribution, and we thank students Helga Helgadóttir and Silja Rúnarsdóttir for their work related to the research. Last but not least we would like to dedicate this research to our co-author, Þórarinn Sveinn Arnarson, who passed away during the submission process.

## Notes

#### Summary of Updates

Several minor changes were made to the text, esp. in Introduction and Discussion, in response to reviewer comments.

