## Supplemental material for "Gas seepage pockmark microbiomes suggest the presence of sedimentary coal seams in the Öxarfjörður graben of NE-Iceland"

Table 9. Description of strains, cultured and isolated on differential and selective media, and conserved in 30% glycerol at -70 °C.

| **Strain no.** | **Media** | **T (°C)** | **Description** | **Strain no.** | **Media** | **T (°C)** | **Description** |
| --- | --- | --- | --- | --- | --- | --- | --- |
| **OX0101** | Naph | 22 | White | **OX0306** | 2,4-D | 22 | Tan |
| **OX0102** | Naph | 22 | White | **OX0307** | 2,4-D | 22 | Greyish white |
| **OX0103** | Naph | 22 | White | **OX0308** | 2,4-D | 22 | White |
| **OX0104** | HEX | 22 | Pink | **OX0309** | 2,4-D | 22 | White |
| **OX0105** | HEX | 22 | Clear-white | **OX0310** | HEX | 22 | Tan |
| **OX0106** | HEX | 22 | Yellow, shiny | **OX0311** | HEX | 22 | Pink |
| **OX0107** | HEX | 22 | White | **OX0312** | HEX | 22 | Pale pink, flat, somewhat swarmy |
| **OX0108** | HEX | 22 | Pink-clear | **OX0313** | HEX | 22 | Yellow |
| **OX0109** | HEX | 22 | Pink | **OX0314** | HEX | 22 | White |
| **OX0110** | HEX | 22 | White, small | **OX0315** | HEX | 22 | Clear-white, swarming |
| **OX0111** | HEX | 22 | Yellow | **OX0316** | Mn | 22 | White with a grey centre |
| **OX0112** | HEX | 22 | Clear-white, swarming | **OX0317** | Mn | 22 | Buff |
| **OX0113** | HEX | 22 | White | **OX0318** | Mn | 22 | Red |
| **OX0116** | Mn | 22 | White with a grey centre | **OX0319** | R2A | 22 | Bright yellow |
| **OX0117** | Mn | 22 | bright yellow | **OX0320** | R2A | 22 | White, irregular |
| **OX0118** | Mn | 22 | Pink | **OX0321** | R2A | 22 | Bright red |
| **OX0119** | Mn | 22 | snowy white | **OX0322** | R2A | 22 | Light yellowish brown, irregular |
| **OX0120** | Mn | 22 | Clear-yellow "fried egg" | **OX0323** | R2A | 22 | Very slightly buff |
| **OX0121** | Mn | 22 | Yellow, shiny | **OX0601** | Naph | 22 | White |
| **OX0122** | R2A | 22 | Bright orange | **OX0602** | Naph | 22 | Orange |
| **OX0123** | R2A | 22 | Bright orange | **OX0603** | Naph | 22 | White, flat |
| **OX0127** | R2A | 22 | Dark yellow | **OX0607** | Naph | 22 | White |
| **Strain no.** | **Media** | **T (°C)** | **Description** | **Strain no.** | **Media** | **T (°C)** | **Description** |
| **OX0128** | R2A | 22 | White w. dark red peak, coherent | **OX0608** | LAS | 22 | Snowy white |
| **OX0129** | R2A | 22 | Bright yellow | **OX0611** | HEX | 22 | White, shiny/oily |
| **OX0130** | R2A | 22 | Pale yellow | **OX0612** | HEX | 22 | White, shiny/oily |
| **OX0620** | Gui | 22 | bright yellow | **OX1215** | Hex | 22°C | Orange |
| **OX0621** | Gui | 22 | White, runny consistency | **OX1216** | BS4 (R2A)* | 15°C | Transparent-white, transparent edge |
| **OX0622** | Gui | 22 | Buff, lobate margins | **OX17B03** | R2A | 22°C | Very transparent |
| **OX0623** | Gui | 22 | bright yellow | **OX17B05** | R2A | 22°C | Light-pink |
| **OX0624** | Gui | 22 | Buff | **OX17B06** | R2A | 22°C | Transparent |
| **OX0625** | R2A | 22 | Bright red | **OX1804** | R2A | 22°C | Red, crawls a lot |
| **OX0626** | R2A | 22 | Pastel yellow | **OX1805** | R2A | 22°C | White-transparent, crawls |
| **OX0627** | R2A | 22 | Yellow "fried egg" | **OX1806** | Hex | 22°C | White, round |
| **OX0628** | R2A | 22 | White | **OX1807** | Hex | 22°C | Light yellow, slimy |
| **OX0629** | R2A | 22 | Clear-white | **OX1808** | Hex | 22°C | Pink |
| **OX0630** | R2A | 22 | Reddish brown | **OX1809** | BS4 (R2A)* | 15° | White-transparent, transparent edge |
| **OX0631** | R2A | 22 | Bright yellow with a clear, flat halo | **OX1905** | R2A | 22°C | Transparent-White, crawls |
| **OX0632** | R2A | 22 | Pink | **OX1906** | R2A | 22°C | White, irregular, crawls |
| **OX0633** | R2A | 22 | Red | **OX1907** | R2A | 22°C | Yellow, lighter edge, crawls |
| **OX1004** | R2A | 22°C | Red, irregular | **OX1908** | Hex | 22°C | White, small, irregular |
| **OX1005** | R2A | 22°C | Small light, crawl much | **OX1909** | Hex | 22°C | Transparent, slimy |
| **OX1006** | R2A | 22°C | Yellow, small, very irregular | **OX2005** | R2A | 22°C | Very light-yellow, crawls, small |
| **OX1007** | R2A | 22°C | Drab (yellow-ish even), irregular | **OX2006** | R2A | 22°C | Light-drab, small, crawls alot |
| **OX1008** | R2A | 22°C | Light-orange, crawls | **OX2007** | R2A | 22°C | Orange, crawls a lot, lighter edge |
| **Strain no.** | **Media** | **T (°C)** | **Description** | **Strain no.** | **Media** | **T (°C)** | **Description** |
| **OX1011** | Hex | 15°C | White | **OX2103** | R2A | 22°C | Light-transparent |
| **OX1012** | BS4 (R2A)* | 35°C | White-transparent, small | **OX2104** | R2A | 22°C | White-transpaeant, crawls a lot |
| **OX1105** | R2A | 22°C | Small, transparent, round | **OX2105** | R2A | 22°C | Yellow, crawls |
| **OX1106** | R2A | 22°C | Small, drab (yellow-ish) | **OX2106** | Hex | 22°C | Orange, round |
| **OX1107** | R2A | 22°C | Light-orange, lighter irregular edge | **OX2107** | Hex | 22°C | Light-orange, round |
| **OX1211** | R2A | 22°C | Yellow, very irregular, lighter edge | **OX1502** | R2A | 22°C | Drab, regular, thin edge |
| **OX1212** | R2A | 22°C | White, round | **OX1503** | R2A | 22°C | Creamy, irregular edge |
| **OX1213** | R2A | 22°C | Orange, very irregular, lighter edge | **OX1504** | R2A | 22°C | Drab, crawls |
| **OX1214** | Hex | 22°C | White, small | **OX1505** | Hex | 15°C | Very light orange, small |
| **OX1602** | R2A | 22°C | White-transparent, thin edge | **SX1204** | R2A | 22°C | Light-tranparent, transparent edge |
| **OX1603** | R2A | 22°C | Yellow, regular, stuck to the agar | **SX1205** | R2A | 22°C | Light-pink, cramped, crawls a lot |
| **OX1604** | Hex | 15°C | White, small | **SX1206** | R2A | 22°C | Light-pink, small |
| **OX1702** | R2A | 22°C | Creamy, transparent, irregular edge | **SX1208** | Hex | 22°C | Light-transparent |
| **OX2208** | R2A | 22°C | Light-orange, crawls | **SX1209** | Hex | 22°C | Light orange, round |
| **OX2209** | Hex | 22°C | White, small, crawls | **SX1210** | Hex | 22°C | Light-white, small |
| **OX2210** | Hex | 22°C | White, small | **SX1212** | Hex | 22°C | White, round |
| **OX2305** | R2A | 22°C | Light-transparent, crawls | **SX1213** | Hex | 22°C | White, very slimy |
| **OX2307** | R2A | 22°C | Orange, lighter edge, irregular | **SX1214** | Hex | 22°C | White-transparent, small |
| **OX2308** | R2A | 22°C | Orange, round | **SX1215** | Hex | 22°C | Light-drab, round |
| **Strain no.** | **Media** | **T (°C)** | **Description** | **Strain no.** | **Media** | **T (°C)** | **Description** |
| **OX2309** | R2A | 22°C | Red-pink, round, lighter edge | **SX201** | R2A | 22°C | White-transparent, crawls a lot |
| **OX2310** | R2A | 22°C | White w/ zone | **SX202** | R2A | 22°C | Light-transparent-yellow, crawls |
| **OX2311** | R2A | 22°C | Orange, shines, crawls | **SX203** | R2A | 22°C | Pink, crawls, slimy |
| **OX2312** | R2A | 22°C | Light-transparent, crawls | **SX204** | R2A | 22°C | Light-pink, big transparent edge |
| **OX2313** | Hex | 22°C | Orange, slimy, transparent | **SX301** | R2A | 22°C | Light-pink, big, slimy |
| **OX2314** | BS4 (R2A)* | 22°C | White-transparent, transparent edge | **SX504** | R2A | 22°C | White, irregular, crawls a lot |
| **OX2404** | R2A | 22°C | Light-yellow, crawls | **SX601** | R2A | 22°C | White, lighter edge, crawls a lot |
| **OX2405** | R2A | 22°C | Light semi-transparent, crawls | **SX602** | R2A | 22°C | White, crawls |
| **OX2406** | Hex | 22°C | Light-orange, irregular | **SX0505** | Hex | 22°C | Pink, round |
| **OX2514** | Hex | 22°C | Light orange | **SX1201** | R2A | 22°C | White, crawls a lot |
| **OX2515** | Hex | 22°C | Orange | **SX1202** | R2A | 22°C | Shiny white, crawls |
| **OX2516** | 9K (R2A)* | 15° | Very light-pink, small, round | **SX0501** | R2A | 22°C | Light-pink, lighter edge, slimy |
| **SX0302** | R2A | 22°C | Light-white, crawls (long branches) | **SX0502** | R2A | 22°C | Pink, crawls |
| **SX0303** | Hex | 22°C | Transparent | **SX0503** | R2A | 22°C | Dark-yellow, crawls a lot, very slimy |
| **SX0304** | Hex | 22°C | White, small , round |  |  |  |  |

Table 10. Taxanomic assignment of strains identified by 16S rDNA sequencing and their GenBanka ccession numbers.

| Strain | EzTaxon assignment | Seq. length | GenBank accession number | | %ID | | Order | Site^1^ | |
| --- | --- | --- | --- | --- | --- | --- | --- | --- | --- |
| **Alphaproteobacteria** | |  |  | |  | |  |  | |
| OX0102 | *Brevundimonas bullata* | 636 | MG576001 | | 99.8 | | Caulobacterales | SX | |
| OX0309 | *Brevundimonas bullata* | 1394 | MG576003 | | 99.9 | | Caulobacterales | SX | |
| OX2008 | *Brevundimonas halotolerans* | 1423 | MG576002 | | 98.9 | | Caulobacterales | AEX | |
| SX0604 | *Porphyrobacter colymbi* | 349 | MG576010 | | 98.6 | | Sphingomonadales | | SX |
| OX0620 | *Sphingobium xenophagum* | 1451 | MG576011 | | 99.4 | | Sphingomonadales | | SX |
| OX0106 | *Rhizobium selenitireducens* | 804 | MG576004 | | 99.9 | | Rhizobiales | SX | |
| OX0119 | *Rhizobium selenitireducens* | 1131 | MG576007 | | 99.9 | | Rhizobiales | SX | |
| OX1214 | *Rhizobium sphaerophysae* | 872 | MG576006 | | 98.1 | | Rhizobiales | AEX | |
| SX1214 | *Rhizobium selenitireducens* | 1447 | MG576005 | | 97.3 | | Rhizobiales | SX | |
| OX1314 | *Paracoccus homiensis* | 721 | MG576009 | | 98.1 | | Rhodobacterales | AEX | |
| OX0632 | *Cereibacter changlensis* | 1396 | MG576008 | | 99.9 | | Rhodobacterales | SX | |
| **Betaproteobacteria** | |  |  | |  | |  |  | |
| OX0105 | *Acidovorax radicis* | 744 | MG576016 | | 98.8 | | Burkholderiales | SX | |
| OX0120 | *Acidovorax radicis* | 1472 | MG576017 | | 99.7 | | Burkholderiales | SX | |
| OX0124 | *Acidovorax radicis* | 1380 | MG576018 | | 99.5 | | Burkholderiales | SX | |
| OX0611 | *Paucibacter toxinivorans* | 1441 | MG576022 | | 97.9 | | Burkholderiales | SX | |
| OX1216 | *Paraburkholderia fungorum* | 1404 | MG576015 | | 99.9 | | Burkholderiales | AEX | |
| OX1313 | *Paraburkholderia fungorum* | 1489 | MG576012 | | 99.9 | | Burkholderiales | AEX | |
| OX1403 | *Paraburkholderia fungorum* | 1492 | MG576013 | | 100 | | Burkholderiales | AEX | |
| OX1702 | *Paraburkholderia fungorum* | 959 | MG576014 | | 100 | | Burkholderiales | AEX | |
| OX0630 | *Rhodoferax fermentans* | 1360 | MG576019 | | 98.5 | | Burkholderiales | AEX | |
| OX0321 | *Rhodoferax saidenbachensis* | 1440 | MG576020 | | 99.3 | | Burkholderiales | AEX | |
| OX0130 | *Variovorax ginsengisoli* | 1465 | MG576021 | | 99.4 | | Burkholderiales | AEX | |
| OX0627 | *Dechloromonas hortensis* | 1501 | MG576023 | | 98.8 | | Rhodocyclales | SX | |
| **Gammaproteobacteria** | |  |  | |  | |  |  | |
| OX0110 | *Aeromonas popoffii* | 1122 | MG576027 | | 99.8 | | Aeromonadales | SX | |
| OX0112 | *Aeromonas popoffii* | 1506 | MG576028 | | 99.7 | | Aeromonadales | SX | |
| OX0612 | *Aeromonas popoffii* | 1451 | MG576029 | | 99.7 | | Aeromonadales | SX | |
| OX0622 | *Aeromonas hydrophila* | 960 | MG576025 | | 100 | | Aeromonadales | SX | |
| SX0303 | *Aeromonas piscicola* | 876 | MG576026 | | 100 | | Aeromonadales | SX | |
| OX0617 | *Aeromonas cavernicola* | 721 | MG576024 | | 100 | | Aeromonadales | SX | |
| OX0631 | *Pseudomonas pictorum* | 1517 | MG576051 | | 99.2 | | Pseudomonadales | SX | |
| OX1807 | *Pseudomonas aestusnigri* | 671 | MG576041 | | 99.4 | | Pseudomonadales | AEX | |
| OX1808 | *Pseudomonas anguillisepticum* | 1536 | MG576042 | | 97.9 | | Pseudomonadales | AEX | |
| OX0306 | *Pseudomonas extremaustralis* | 1429 | MG576043 | | 99.7 | | Pseudomonadales | AEX | |
| OX0322 | *Pseudomonas extremaustralis* | 1342 | MG576044 | | 99.7 | | Pseudomonadales | AEX | |
| OX1110 | *Pseudomonas guineae* | 1502 | MG576045 | | 99.8 | | Pseudomonadales | AEX | |
| OX0613 | *Pseudomonas linyingensis* | 1450 | MG576046 | | 99.9 | | Pseudomonadales | AEX | |
| SX0305 | *Pseudomonas mandelii* | 1416 | MG576047 | | 99.9 | | Pseudomonadales | SX | |
| SX0307 | *Pseudomonas mandelii* | 1503 | MG576048 | | 99.9 | | Pseudomonadales | SX | |
| SX1216 | *Pseudomonas mandelii* | 1503 | MG576049 | | 99.9 | | Pseudomonadales | SX | |
| SX0304 | *Pseudomonas peli* | 702 | MG576050 | | 99.5 | | Pseudomonadales | SX | |
| SX1213 | *Pseudomonas vancouverensis* | 1500 | MG576052 | | 99.7 | | Pseudomonadales | SX | |
| OX0304 | *Pseudomonas veronii* | 702 | MG576053 | | 99.6 | | Pseudomonadales | SX | |
| OX0601 | *Acinetobacter pakistanensis* | 1501 | MG576039 | | 100 | | Pseudomonadales | SX | |
| OX0621 | *Acinetobacter pakistanensis* | 839 | MG576040 | | 100 | | Pseudomonadales | SX | |
| OX0619 | *Shewanella putrefaciens* | 1511 | MG576030 | | 99.5 | | Alteromonadales | SX | |
| OX0103 | *Rahnella aquatilis* | 576 | MG576034 | | **94.4** | | Enterobacteriales | SX | |
| OX0101 | *Rahnella inusitata* | 1507 | MG576035 | | 99.7 | | Enterobacteriales | SX | |
| OX0604 | *Shigella flexneri* | 1505 | MG576036 | | 99.7 | | Enterobacteriales | SX | |
| OX0606 | *Shigella flexneri* | 1510 | MG576037 | | 99.7 | | Enterobacteriales | SX | |
| OX0607 | *Escherichia coli* | 858 | MG576038 | | 99.5 | | Enterobacteriales | SX | |
| OX1012 | *Escerichia fergusonii* | 1504 | MG576033 | | 99.7 | | Enterobacteriales | AEX | |
| OX1208 | *Rheinheimera soli* | 1515 | MG576032 | | 99.0 | | Chromatiales | AEX | |
| OX1909 | *Rheinheimera aestuari* | 977 | MG576031 | | 99.1 | | Chromatiales | AEX | |
| **Bacilli** |  |  |  | |  | |  |  | |
| OX0301 | *Paenibacillus xylanexedens* | 1214 | MG575992 | | 98.8 | | Bacillales | SX | |
| OX0302 | *Paenibacillus xylanexedens* | 1510 | MG575993 | | 99.7 | | Bacillales | SX | |
| OX2310 | *Paenibacillus tundrae* | 1497 | MG575991 | | 99.3 | | Bacillales | AEX | |
| OX0307 | *Paenibacillus terrae* | 1502 | MG575990 | | 98.9 | | Bacillales | SX | |
| OX2205 | *Paenibacillus alba* | 961 | MG575989 | | 99.7 | | Bacillales | AEX | |
| OX0310 | *Jeotgalibacillus campisalis* | 1484 | MG575994 | | 99.4 | | Bacillales | SX | |
| OX0626 | *Staphylococcus argentus* | 1409 | MG575995 | | 99.9 | | Bacillales | SX | |
| OX0317 | *Psychrobacillus psychrodurans* | 1496 | MG575988 | | 99.5 | | Bacillales | SX | |
| OX1213 | *Bacillus aquimaris* | 1507 | MG575973 | | 99.5 | | Bacillales | AEX | |
| OX2513 | *Bacillus halmapalus* | 872 | MG575974 | | 98.8 | | Bacillales | AEX | |
| OX1505 | *Bacillus hwajinpoensis* | 1262 | MG575975 | | 99.0 | | Bacillales | AEX | |
| OX2509 | *Bacillus hwajinpoensis* | 1282 | MG575976 | | 99.4 | | Bacillales | AEX | |
| OX1011 | *Bacillus oceanisediminis* | 937 | MG575977 | | 100 | | Bacillales | AEX | |
| OX1604 | *Bacillus oceanisediminis* | 363 | MG575978 | | 99.2 | | Bacillales | AEX | |
| OX1805 | *Bacillus safensis* | 885 | MG575979 | | 99.6 | | Bacillales | AEX | |
| OX1004 | *Bacillus vietnamensis* | 1434 | MG575980 | | 99.9 | | Bacillales | AEX | |
| OX1006 | *Bacillus vietnamensis* | 1351 | MG575981 | | 99.9 | | Bacillales | AEX | |
| OX1007 | *Bacillus vietnamensis* | 1508 | MG575982 | | 99.9 | | Bacillales | AEX | |
| OX1210 | *Bacillus vietnamensis* | 1394 | MG575983 | | 99.8 | | Bacillales | AEX | |
| OX1212 | *Bacillus vietnamensis* | 1509 | MG575984 | | 99.9 | | Bacillales | AEX | |
| OX2308 | *Bacillus vietnamensis* | 1205 | MG575985 | | 99.8 | | Bacillales | AEX | |
| OX2309 | *Bacillus vietnamensis* | 1059 | MG575986 | | 99.7 | | Bacillales | AEX | |
| OX2506 | *Bacillus vietnamensis* | 1374 | MG575987 | | 99.9 | | Bacillales | AEX | |
| OX2515 | *Exiguobacterium oxidotolerans* | 861 | MG575998 | | 99.8 | | Lactobacillales | AEX | |
| OX2106 | *Exiguobacterium profundum* | 754 | MG575999 | | 99.9 | | Lactobacillales | AEX | |
| OX2313 | *Exiguobacterium profundum* | 1053 | MG576000 | | 99.4 | | Lactobacillales | AEX | |
| OX1107 | *Exiguobacterium profundum* | 809 | MG575996 | | 99.5 | | Lactobacillales | AEX | |
| OX1109 | *Exiguobacterium profundum* | 1463 | MG575997 | | 98.2 | | Lactobacillales | AEX | |
| **Flavobacteria** | |  |  | |  | |  |  | |
| OX0623 | *Flavobacterium glaciei* | 992 | MG575964 | | 99.0 | | Flavobacteriales | SX | |
| OX0125 | *Flavobacterium granuli* | 1446 | MG575965 | | 97.8 | | Flavobacteriales | SX | |
| OX0129 | *Flavobacterium granuli* | 1449 | MG575966 | | 97.8 | | Flavobacteriales | SX | |
| OX0311 | *Flavobacterium hydatis* | 1506 | MG575967 | | 97.6 | | Flavobacteriales | SX | |
| OX0314 | *Flavobacterium succinians* | 1496 | MG575970 | | 98.4 | | Flavobacteriales | SX | |
| OX0122 | *Flavobacterium succinians* | 1451 | MG575968 | | 98.5 | | Flavobacteriales | SX | |
| OX0123 | *Flavobacterium succinians* | 1402 | MG575969 | | 98.4 | | Flavobacteriales | SX | |
| OX0127 | *Flavobacterium terrigena* | 1447 | MG575971 | | 98.6 | | Flavobacteriales | SX | |
| **Cytophagia** | |  |  | |  | |  |  | |
| OX0104 | *Algoriphagus alkaliphilus* | 1481 | MG575959 | | 97.9 | | Cytophagales | SX | |
| OX0118 | *Algoriphagus alkaliphilus* | 1471 | MG575960 | | 97.5 | | Cytophagales | SX | |
| OX0108 | *Aquiflexum balticum* | 1060 | MG575961 | | **94.7** | | Cytophagales | SX | |
| OX0126 | *Aquiflexum balticum* | 1459 | MG575962 | | **95.5** | | Cytophagales | SX | |
| OX0312 | *Aquiflexum balticum* | 1459 | MG575963 | | **95.5** | | Cytophagales | SX | |
| **Sphingobacteria** | |  |  | |  | |  |  | |
| OX0625 | *Pedobacter ruber* | 1452 | MG575972 | 98.5 | | Sphingobacteriales | | SX | |
| **Actinobacteria** | |  |  | |  | |  |  | |
| OX0615 | *Oerskovia paurometabola* | 1353 | MG575950 | | 98.4 | | Micrococcales | SX | |
| OX0315 | *Cryobacterium arcticum* | 1463 | MG575951 | | 99.3 | | Micrococcales | SX | |
| OX0319 | *Cryobacterium arcticum* | 932 | MG575952 | | 100 | | Micrococcales | SX | |
| OX0117 | *Arthrobacter alpinus* | 1471 | MG575953 | | 99.0 | | Micrococcales | SX | |
| OX0308 | *Arthrobacter humicola* | 1460 | MG575954 | | 99.9 | | Micrococcales | SX | |
| OX0313 | *Arthrobacter oryzae* | 1494 | MG575955 | | 99.3 | | Micrococcales | SX | |
| OX0107 | *Pseudarthrobacter siccitolerans* | 1487 | MG575956 | | 99.0 | | Micrococcales | SX | |
| OX0614 | *Sanguibacter suarezii* | 1243 | MG575957 | | 99.1 | | Micrococcales | SX | |
| OX0316 | *Streptomyces clavifer* | 1156 | MG575958 | | 99.6 | | Streptomycetales | SX | |
| SX1205 | *Rhodococcus globerulus* | 1294 | MG575949 | | 99.7 | | Corynebacteriales | SX | |
| SX0206 | *Rhodococcus globerulus* | 1482 | MG575948 | | 99.7 | | Corynebacteriales | SX | |

^1^ *Site SX is at Skógaeyralón (66.15°N, 16.62°W). Vegetated, wetland area. Some suspected gas seepage, but little or no evidence of geothermal influence. Site AER is at Skógakíll near Ærlækjarsel (66.17°N, 16.57°W). Mostly barren, sandy, estuarine area characterized by geothermal activity.*
